# Targeting Microbial Bile Salt Hydrolase Reprograms Bile Acid Metabolism and Ameliorates Metabolic Dysfunction–Associated Steatohepatitis in Mice

**DOI:** 10.64898/2026.05.12.724693

**Authors:** Wenchao Wei, Radka Graf, Yanhan Wang, Christopher J. Oalmann, Jennifer T. Lau, Xinyi Wang, Melanie Chien, Mary C. Conrad, Jonah Simon, Souradipta Ganguly, Tomoo Yamazaki, Aenne Harberts, Sainan Chen, Marcos Fernandez Fondevila, Debanjan Dhar, Stewart A. Campbell, Rebecca K. Senter, Bernd Schnabl

## Abstract

Microbial bile salt hydrolase (BSH) plays a central role in shaping bile acid composition and gut–liver metabolic signaling, yet its therapeutic potential in metabolic dysfunction–associated steatohepatitis (MASH) remains incompletely defined. Here, we evaluated the efficacy of the non-absorbable BSH inhibitor GR-7 in a diet induced mouse model of steatohepatitis using early and late intervention strategies with different dosing regimens. GR-7 reduced food intake and exerted stage- and dose-dependent therapeutic effects, with early intervention robustly suppressing hepatic fibrosis even at low dose, whereas late-stage administration of high-dose GR-7 markedly reduced hepatic steatosis and inflammation, as evidenced by decreased liver weight, hepatic triglyceride and cholesterol levels, and plasma ALT. Although late intervention did not result in statistically significant histological reversal of fibrosis, a trend toward improvement was observed, together with suppression of fibrogenic gene expression, suggesting that prolonged treatment may further enhance antifibrotic efficacy. Mechanistically, GR-7 effectively inhibited microbial BSH activity in vivo, leading to reduced cecal unconjugated primary and secondary bile acids—including deoxycholic acid and lithocholic acid, which was associated with improved gut barrier integrity and reduced hepatic inflammation. In parallel, BSH inhibition reprogrammed hepatic bile acid metabolism toward activation of the alternative CYP27A1-mediated synthesis pathway, accompanied by reduced food intake, thereby contributing to improved hepatic lipid accumulation. Furthermore, late-stage high-dose treatment selectively remodeled the hepatic immune landscape rather than fully restoring homeostasis, highlighting immune recalibration as a key component of therapeutic response. Together, these findings identify microbial BSH inhibition as a promising microbiome-targeted therapeutic strategy for MASH.

**Highlights:**

- The non-absorbable BSH inhibitor GR-7 improves steatosis, inflammation, and fibrosis in of Western diet-induced steatohepatitis model in mice in a dose-dependent manner.
- GR-7 reduces food intake and body weight gain.
- GR-7 reduces cytotoxic secondary bile acids, including DCA and LCA.
- GR-7 reprograms hepatic bile acid metabolism and immune responses.

## Introduction

Metabolic dysfunction–associated steatotic liver disease (MASLD) is an increasingly prevalent metabolic disorder, affecting up to 38% of the adult population globally.^1^ Its progressive form, metabolic dysfunction–associated steatohepatitis (MASH), is characterized by hepatocellular injury and inflammation, and represents a critical disease stage that predisposes patients to fibrosis, cirrhosis, and hepatocellular carcinoma.^2^ Consequently, MASLD/MASH are associated with increased liver-related morbidity and mortality and are a major driver of the rising need for liver transplantation.^3^ These significant clinical and societal burdens show the urgent need for effective pharmacological therapies for MASH. In 2024, resmetirom became the first drug approved by the U.S. Food and Drug Administration for the treatment of MASH.^4^ As a selective thyroid hormone receptor-β agonist, resmetirom reduces hepatic lipid accumulation and improves liver inflammation as well as moderate to advanced fibrosis, representing a major milestone in MASH therapeutics.^5^ In addition, semaglutide, an incretin-based therapy, has also demonstrated therapeutic efficacy in MASH, including improvements in steatohepatitis and fibrosis-related endpoints.^6^ Nevertheless, despite these advances, there remains a substantial unmet need for additional effective therapies targeting MASH.

Bile acid signaling has emerged as a key regulator of metabolic and inflammatory pathways through the gut–liver axis and a promising therapeutic target for MASLD/MASH.^7–9^ Primary bile acids synthesized from cholesterol in the liver are conjugated and secreted into the intestine to facilitate lipid absorption, where they are deconjugated by microbial bile salt hydrolase (BSH) and subsequently converted into secondary bile acids.^10^ Previous clinical studies have shown that BSH activity is highly associated with MASLD prevalence and disease severity.^11–13^ Moreover, experimental evidence indicates that modulation of BSH activity in a single bacterial strain can markedly alter host bile acid composition and systemic metabolic homeostasis.^14^ Notably, inhibition of microbial BSH improves gut barrier function and attenuates liver injury, underscoring its therapeutic potential in liver disease.^15^

In this study, we assessed the therapeutic efficacy of GR-7, a bile salt hydrolase (BSH) inhibitor, in mouse models of MASH. GR-7 has been previously described as an α-fluoromethyl ketone–based inhibitor that covalently targets the catalytic cysteine residue of BSH.^15^

## Materials and Methods

### Metabolic Dysfunction–Associated Steatohepatitis (MASH) mouse model

All animal experiments were performed in at least two independent batches. Male C57BL/6 mice deficient in the *Alms1* gene (Foz/Foz), a hyperphagic model of metabolic dysfunction–associated steatohepatitis (MASH), were used as described by us.^16^ At 6 weeks of age, mice were fed a Western diet (WD) for 24 weeks to induce MASH. The WD provided 40% kcal from fat (milk fat, 12% saturated) and 0.2% cholesterol and was enriched in fructose (AIN-76A Western Diet 5342, Catalog #TD.200289, Envigo). The bile salt hydrolase (BSH) inhibitor GR-7 was incorporated into the diet by the manufacturer at concentrations of 0.034% (low dose) or 0.1% (high dose), and all diets were irradiated. Mice were randomly assigned to six groups: chow diet (n=7), WD alone (n=12), WD + low-dose GR-7 early intervention (n=10), WD + high-dose GR-7 early intervention (n=12), WD + low-dose GR-7 late intervention (n=6), and WD + high-dose GR-7 late intervention (n=12) (Figure 1). In the early intervention model, GR-7 was introduced after 3 weeks of WD feeding, whereas in the late intervention, treatment began after 12 weeks of WD feeding. Chow- and WD-fed animals served as shared controls for both intervention models. One mouse in the WD group died between week 6 and 7, two mice in the WD plus low-dose GR-7 early-intervention died between weeks 21 and 23, and three mice in the WD plus high-dose GR-7 late-intervention group died between weeks 2 and 5 prior to initiation of GR-7 treatment and while receiving WD alone. Tissue samples were not collected from these six mice. Mouse body weight and food intake were recorded weekly. Food intake was calculated on a per-mouse, per-day basis from cage-level consumption. All mice were euthanized at week 24 following an 8-hour fast.

**Figure 1.**
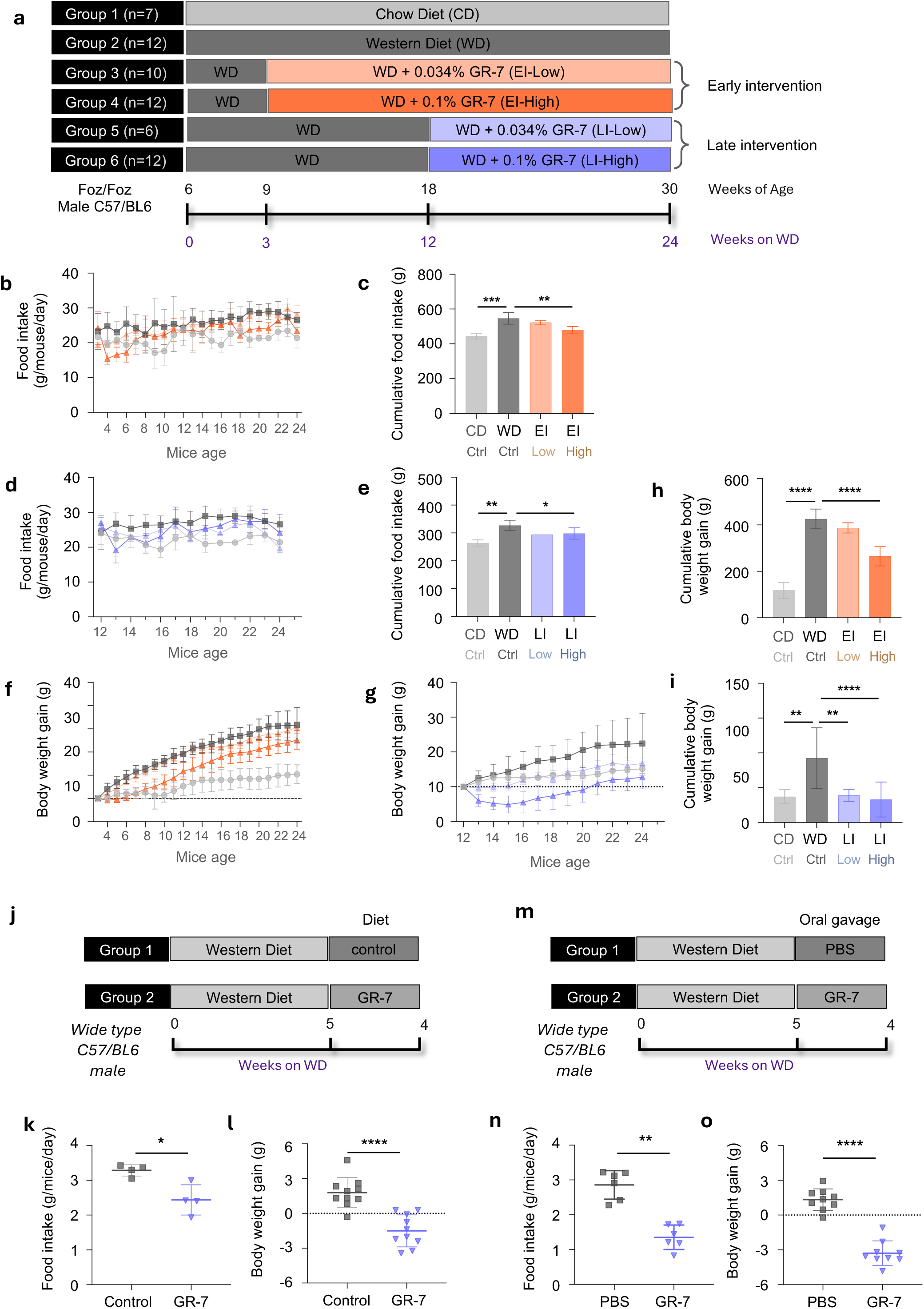
The BSH inhibitor GR-7 reduces food intake and body weight gain in western diet–fed mice. **a**, Experimental design. Male C57BL/6 *Foz/Foz* mice were fed chow diet (CD) or western diet (WD). GR-7 was administered either as early intervention (EI) starting at week 3 of WD feeding or as late intervention (LI) starting at week 12, at low (0.034%) or high (0.1%) dietary doses. Group sizes are indicated. **b**, Longitudinal food intake during early intervention. **c**, Cumulative food intake during early intervention. **d**, Longitudinal food intake during late intervention. **e**, Cumulative food intake during late intervention. **f**, Longitudinal body weight gain during early intervention. **g**, Longitudinal body weight gain during late intervention. **h**, Cumulative body weight gain during early intervention. **i**, Cumulative body weight gain during late intervention. **j**, Short-term dietary GR-7 intervention schematic in WD-fed wild-type mice. **k**, Daily food intake following short-term dietary GR-7 treatment. **l**, Body weight gain during short-term dietary GR-7 treatment. **m**, Oral gavage intervention schematic in WD-fed wild-type mice receiving PBS or GR-7. **n**, Daily food intake following oral gavage of PBS or GR-7. **o**, Body weight gain following oral gavage of PBS or GR-7. Data are shown as mean ± SD. Each symbol represents one mouse/one cage. Statistical significance was determined by one-way ANOVA with Bonferroni’s multiple-comparisons test. For comparisons between two groups, Mann–Whitney U test was used. * *P* < 0.05; ** *P* < 0.01; *** *P* < 0.001; **** *P* < 0.0001.

### One-week GR-7 treatment study to assess food intake and body weight gain (dietary administration)

Wild-type male C57BL/6 mice (6 weeks of age) were obtained from Charles River Laboratories and fed a WD for 5 weeks. Mice were then randomized into two groups and treated for an additional 1 week with either WD alone (n = 10) or WD supplemented with high-dose GR-7 (0.1%, w/w; n = 10). Food intake and body weight were monitored throughout the treatment period, and body weight gain was calculated. Food intake was calculated on a per-mouse, per-day basis from cage-level consumption.

### One-week GR-7 treatment study to assess food intake and body weight gain (oral gavage)

Wild-type male C57BL/6 mice (6 weeks of age) were obtained from Charles River Laboratories and fed a WD for 5 weeks. Mice were then maintained on WD and treated once daily for 1 week with either PBS (n = 9) or GR-7 (77 mg in 100 µL PBS; n = 9) by oral gavage. The GR-7 dose was matched to the high-dose regimen used during late intervention based on average food intake. Food intake and body weight were recorded daily, and body weight gain was calculated over the treatment period. Food intake was calculated on a per-mouse, per-day basis from cage-level consumption.

### Animal housing and ethics

Mice were housed under standard conditions with a 12 h light/12 h dark cycle, controlled temperature and humidity, and ad libitum access to food and water. All animal procedures were approved by the Institutional Animal Care and Use Committee at the University of California, San Diego (protocol number S09042) and conducted in accordance with institutional and ARRIVE guidelines.

### RNA extraction and quantitative real-time PCR

Total RNA was extracted from intestinal tissues using TRIzol reagent (Invitrogen) according to the manufacturer’s instructions. Genomic DNA contamination was removed using the DNA-free™ DNA Removal Kit (Ambion), and complementary DNA (cDNA) was synthesized using the High-Capacity cDNA Reverse Transcription Kit (Applied Biosystems). Primer sequences for mouse genes were originally obtained from the NIH qPrimerDepot database. Gene expression was quantified using SYBR Green chemistry (Bio-Rad Laboratories) on an ABI StepOnePlus Real-Time PCR System. Relative mRNA expression in the ileum was calculated using the ΔΔCt method and normalized to Gapdh as the internal control. The primer sequences were as follows: mouse *Gapdh* (forward: 5′-TTGATGGCAACAATCTCCAC-3′; reverse: 5′- CGTCCCGTAGACAAAATGGT-3′) and mouse *Fxr* (*Nr1h4*) (forward: 5′-GGCAGAATCTGGATTTGGAATCG-3′; reverse: 5′-GCCCAGGTTGGAATAGTAAGACG-3′).

### Histological staining and image analysis

Formalin-fixed tissue samples were embedded in paraffin (Paraplast Plus, McCormick), sectioned at 4 µm, and stained with hematoxylin and eosin (H&E; Surgipath) or 0.1% picrosirius red (PSR; Color Index 35780, 365548; Sigma-Aldrich). PSR-stained fibrotic area was quantified using ImageJ software (version 1.53k; Java 13.0.6). For detection of neutral lipid accumulation, liver tissues were embedded in OCT compound, sectioned at 10 µm on a cryostat, and stained with Oil Red O (Sigma-Aldrich). Quantification of Oil Red O–positive area was performed using ImageJ.

### Biochemical analysis

Plasma alanine aminotransferase (ALT) levels were measured using the ALT (SGPT) Kinetic Liquid Assay (Teco Diagnostics). Hepatic triglyceride and cholesterol concentrations were quantified using Triglyceride Liquid Reagents and Cholesterol Liquid Reagents kits, respectively (Pointe Scientific). Plasma lipopolysaccharide (LPS) levels were assessed by enzyme-linked immunosorbent assay (ELISA; Lifeome Biolabs). Liver hydroxyproline content was quantified from 100 mg of liver tissue. Samples were homogenized in 6 N HCl (3750-32; USABlueBook) using lysing matrix C tubes and the Mini-BeadBeater-96, followed by incubation at 110 °C for 24 hours. The lysates were then filtered using Whatman grade 595 1/2 filter paper (WHA10311644; Sigma-Aldrich). After reaction with chloramine T (C9887; Sigma-Aldrich) and Ehrlich’s perchloric acid solution (AC168760250; Thermo Fisher Scientific), absorbance was measured at 558 nm using a VersaMax microplate reader (Molecular Devices LLC, Sunnyvale, CA).

### Bile acid extraction and quantification

Cecum samples were weighed and homogenized in water with metal beads, followed by protein precipitation with acetonitrile. Supernatants were diluted with a mixture of internal standard solution containing 14 deuterium-labeled bile acids and acetonitrile prior to analysis. Bile acids were quantified by UPLC-MS/MS using an Agilent 1290 UHPLC coupled to an Agilent 6495B triple quadrupole mass spectrometer operated in multiple-reaction monitoring mode with negative-ion detection (Creative Proteomics, Inc., Shirley, NY). Separation was performed on a Waters C18 column (2.1 × 150 mm, 1.7 µm) with a binary solvent gradient of 0.01% formic acid in water and acetonitrile. Calibration curves were constructed from serially diluted bile acid standards (0.00002–10 µM), and sample concentrations were determined using analyte-to-internal standard peak area ratios.

### RNA seq and data analysis

Total RNA was extracted from liver tissues using the QIAGEN RNeasy Mini Tissue Extraction Kit, with genomic DNA removed via DNase treatment. RNA quality and quantity were assessed prior to library preparation. Libraries were constructed from 100 ng of RNA per sample using the QIAGEN QIAseq UPXome RNA Library Kit with QIAseq FastSelect rRNA depletion and N6-T reverse transcription primers. After reverse transcription, 12 samples were pooled per batch and amplified for 15 PCR cycles. Sequencing was performed on the MGI DNBSEQ-T7 platform. Raw reads were quality-checked, trimmed, and aligned to the reference genome using standard RNA-seq pipelines (Gnomix, South Australia, Australia). Gene-level expression data were imported into R (v4.5.2), merged across samples by gene identifier, and missing values were set to zero. Genes with positive expression in at least one-third of samples were retained to reduce low-abundance noise, yielding 15,519 high-quality genes. Expression consistency across samples was assessed using log10-transformed TPM distributions and quality control metrics. Differential expression analysis was performed on protein-coding exon count matrices using DESeq2 after low-expression filtering (CPM > 1 in at least three samples), followed by size-factor normalization and variance-stabilizing transformation. Differential expression was modeled using a negative binomial generalized linear model with experimental condition as the design factor, and significant genes were defined by adjusted P < 0.05 and contrast-specific fold-change thresholds. KEGG pathway enrichment analysis was conducted on significantly upregulated and downregulated genes (adjusted P < 0.1, |fold change| ≥ 1.2), with enriched pathways ranked by adjusted P value and fold enrichment and visualized using dot plots.

### Immune cell isolation, staining, and Mass cytometry (CyTOF) Analysis

Single-cell suspensions were prepared from freshly isolated mouse livers by mechanical dissociation, red blood cell lysis, and Percoll density-gradient centrifugation. Purified cells were washed, resuspended in complete medium, and counted prior to analysis. For mass cytometry, cells were stained with a panel of metal-conjugated antibodies. Briefly, cells were first labeled with cisplatin (Cell-ID cisplatin, Fluidigm) for live/dead discrimination, followed by sample barcoding (Cell-ID 20-Plex Pd Barcoding kit, PN PRD023, Fluidigm), and staining of surface markers. Cells were then fixed, permeabilized, and stained for intracellular cytokines and cytoplasmic proteins. After final fixation, DNA was labeled with an intercalator (Cell-ID Intercalator, Fluidigm) prior to acquisition on a CyTOF mass cytometer. The antibody panel included markers defining major immune lineages, activation and exhaustion states, and effector cytokines (see Supplemental Table1 for details). Samples were resuspended in Cell Acquisition Solution supplemented with EQ calibration beads and acquired on a CyTOF mass cytometer (Fluidigm), with approximately 300,000–800,000 events collected per sample depending on cell yield. Data were normalized using EQ bead-based signal correction, and quality control steps included removal of beads, doublets, debris, and dead cells based on cisplatin and DNA intercalator signals. Immune populations were defined by sequential gating of CD45^+^ leukocytes and lineage-specific markers using Cytobank and downstream computational analysis workflows (Standard Biotools, Ontario, Canada).

### Statistical analysis

Results are expressed as mean ± standard deviation (SD). At least two technical replicates were performed in each of the groups. Significance was evaluated using One-way analysis of variance (ANOVA) followed by Bonferroni’s multiple comparisons test for experiments involving more than three groups. The mean of each column was compared to the mean of western diet-fed control mice. Statistical comparison between two groups was conducted using a two-tailed unpaired Mann–Whitney U test. A *P* value < 0.05 was considered to be statistically significant. All statistical analyses were performed using GraphPad Prism v10.3.0.

## Results

### GR-7 reduces food intake and attenuates body weight gain in MASH mouse models

To evaluate the efficacy of the BSH inhibitor GR-7 in MASH, we used an established diet-induced MASH model by feeding male *Alms1*-deficient C57BL/6 (*Foz/Foz*) mice a Western diet (WD) for 24 weeks (Fig. 1a).^16^ Chow diet (CD) fed mice served as normal controls. To assess early intervention, mice were administered either a low dose (0.034% weight of drug/weight of chow (w/w)) or a high dose (0.1% w/w) of GR-7 starting after 3 weeks of WD feeding and continued until week 24. To evaluate late intervention, separate cohorts received the same low or high doses of GR-7 beginning after 12 weeks of WD feeding and continued until week 24. Food intake and body weight were monitored weekly throughout the study.

Compared with WD-fed controls, mice receiving the high dose of GR-7 exhibited a significant reduction in food intake during both early and late intervention phases, whereas the low dose of GR-7 showed a nonsignificant downward trend (Figs. 1b–e). High-dose GR-7 significantly attenuated WD-induced weight gain in both early and late intervention settings, while the low dose significantly reduced body weight gain only during the late intervention phase (Figs. 1f–i).

Because *Foz/Foz* mice exhibit hyperphagia, we next sought to determine whether GR-7 exerted similar effects in wild-type mice. A short-term experiment was conducted in male wild-type C57BL/6 mice, which were fed WD for 5 weeks followed by one week of either control diet or WD supplemented with 0.1% GR-7 (Fig. 1j). One week of high-dose GR-7 supplementation significantly reduced both food intake and body weight gain compared with controls (Figs. 1k–l).

To further exclude the possibility that reduced food intake resulted from altered diet palatability or odor, GR-7 administration was switched from dietary supplementation to oral gavage (Fig. 1m). Consistently, mice receiving GR-7 by gavage displayed significantly reduced food intake and body weight gain compared with PBS-treated controls (Figs. 1n–o).

Collectively, these results demonstrate that GR-7 reduces food intake and attenuates body weight gain in both WD-fed *Foz/Foz* and wild-type mice, with greater efficacy observed at higher dosage, supporting a dose-dependent metabolic effect independent of dietary palatability.

### Early intervention with GR-7 attenuates hepatic cholesterol accumulation and fibrosis

To evaluate the efficacy of early intervention with GR-7, a comprehensive assessment of hepatic metabolic and histological parameters was performed. Compared with chow-fed controls, WD feeding resulted in significant increases in liver weight and liver-to-body weight ratio (Figs. 2a–b; all *P* < 0.0001), accompanied by marked elevations in hepatic triglyceride and cholesterol levels and enhanced Oil Red O staining, indicative of substantial hepatic lipid accumulation (Figs. 2c–e, j; all *P* < 0.0001). Early intervention with high-dose GR-7 significantly reduced liver weight, liver-to-body weight ratio, and total hepatic cholesterol compared with WD-fed mice (Figs. 2a, b, d; all *P* < 0.01). Early intervention resulted in lower total hepatic triglycerides and Oil Red O staining, although this was not statistically significant (Figs. 2c, e, j).

**Figure 2.**
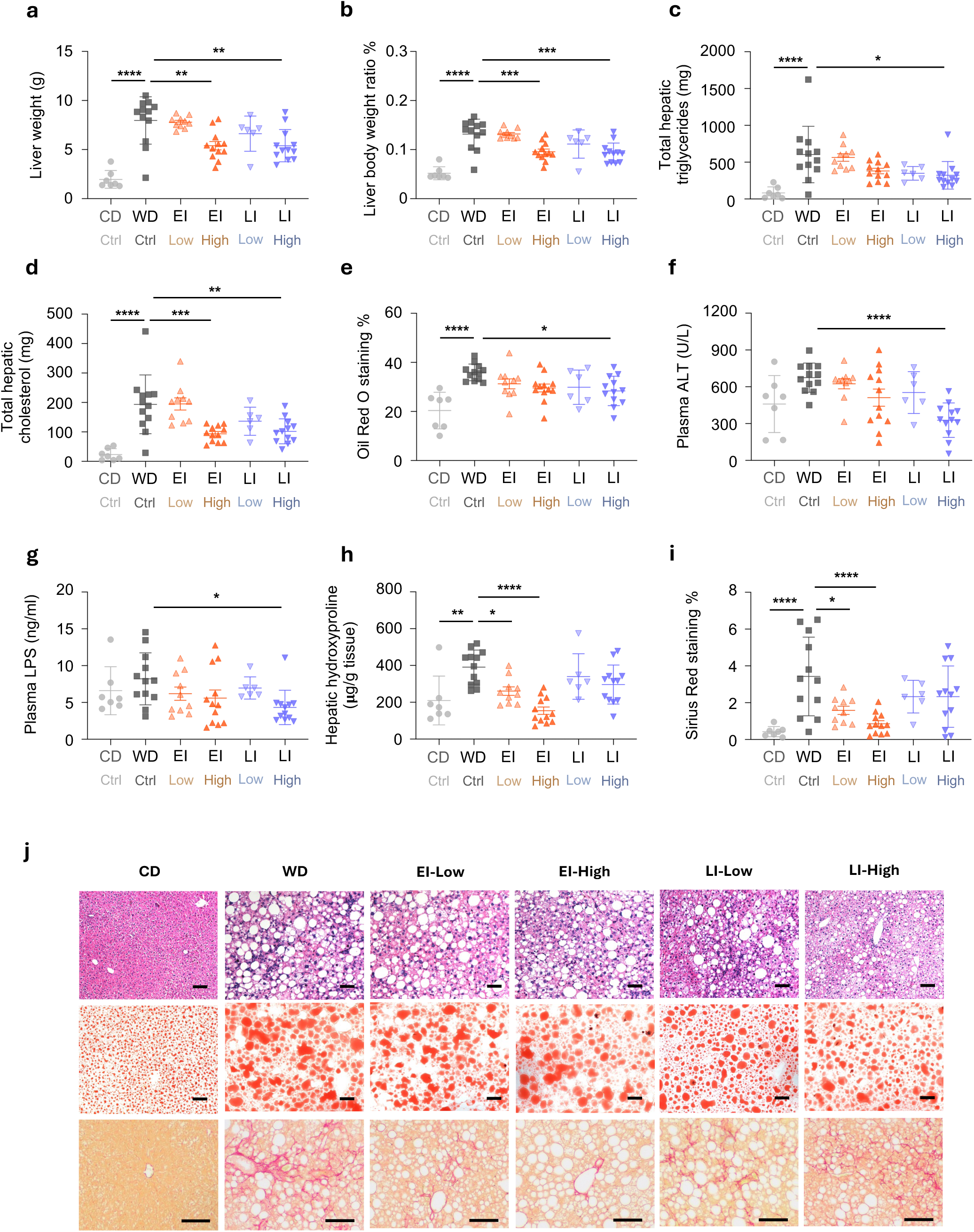
GR-7 ameliorates hepatic steatosis, inflammation, and fibrosis in western diet–fed mice. **a**, Liver weight. **b**, Liver-to-body weight ratio. **c**, Total hepatic triglyceride content. **d**, Total hepatic cholesterol content. **e**, Quantification of Oil Red O–positive area. **f**, Plasma alanine aminotransferase (ALT) levels. **g**, Plasma lipopolysaccharide (LPS) levels. **h**, Hepatic hydroxyproline content. **i**, Quantification of Sirius Red–positive area. **j**, Representative histological images of liver sections from chow diet (CD), western diet (WD), early-intervention low-dose (EI-Low), early-intervention high-dose (EI-High), late-intervention low-dose (LI-Low), and late-intervention high-dose (LI-High) groups. Top row: H&E staining; middle row: Oil Red O staining; bottom row: Sirius Red staining. Scale bars, 50 μm. Data are shown as mean ± SD. Each symbol represents one mouse. Statistical significance was determined by one-way ANOVA with Bonferroni’s multiple-comparisons test. * *P* < 0.05; ** *P* < 0.01; *** *P* < 0.001; **** *P* < 0.0001.

Plasma alanine aminotransferase (ALT) and lipopolysaccharide (LPS) levels were not significantly altered by WD feeding (Figs. 2f–g), likely reflecting elevated baseline values in obese chow-fed *Foz/Foz* mice. In contrast, WD feeding markedly increased hepatic hydroxyproline content and Sirius Red–positive staining, indicating enhanced liver fibrosis (Figs. 2h–j). Notably, administration of GR-7 during early intervention significantly attenuated these fibrosis indices, even at a low dose. Early intervention lowered plasma ALT and LPS levels, although this was not statistically significant (Figs. 2f–g).

Collectively, these findings demonstrate that early intervention with GR-7 confers protection against MASH development, characterized by lower lipid accumulation and robust anti-fibrotic effects even at low doses.

### Late intervention with GR-7 attenuates hepatic lipid accumulation and inflammation

To evaluate the therapeutic efficacy of late intervention with GR-7 in established MASH, hepatic metabolic and histological parameters were assessed. Compared with WD–fed controls, only high-dose GR-7 treatment significantly reduced liver weight, liver-to-body weight ratio, and total hepatic triglyceride and cholesterol levels, findings that were consistent with quantitative Oil Red O staining (Figs. 2a–e, j; all *P* < 0.05). These results indicate that high-dose GR-7 effectively attenuates hepatic lipid accumulation when administered after disease establishment. In addition, therapeutic high-dose GR-7 significantly decreased plasma ALT and LPS levels, suggesting reduced hepatic inflammation and improved intestinal barrier integrity (Figs. 2f–g; all *P* < 0.05). Hepatic fibrosis was lower following late intervention, although this was not significantly different, indicating that the anti-fibrotic effects of GR-7 may require longer treatment (Figs. 2h–j).

Collectively, these findings demonstrate that late intervention GR-7 administration reverses hepatic steatosis and inflammation and improves gut barrier function in a dose-dependent manner.

### Pharmacological inhibition of BSH activity modulates cecal bile acid profiles

To assess the in vivo efficacy of the BSH inhibitor GR-7, targeted bile acid profiling was performed in cecal contents. Unconjugated bile acids constituted more than 90% of the total bile acid in the cecum, consistent with efficient ileal reabsorption of conjugated bile acids and extensive bacterial BSH activity in the cecum. Compared with chow-fed controls, WD feeding markedly increased total primary unconjugated bile acids (Fig. 3a, *P* < 0.0001), whereas total primary conjugated bile acids were not significantly altered (Fig. 3b). This diet-induced enhancement of bile acid deconjugation was significantly reversed by high-dose GR-7 treatment in both early and late intervention settings, with pronounced reductions in α-muricholic acid (α-MCA) and β-muricholic acid (β-MCA) (Fig. 3a, all *P* < 0.05), consistent with effective inhibition of bacterial BSH activity in vivo. There was a trend toward lower levels of cholic acid (CA) and chenodeoxycholic acid (CDCA) with high-dose GR-7 treatment. WD feeding also significantly increased total secondary bile acids, including both conjugated and unconjugated species (Figs. 3c–d, all *P* < 0.001), reflecting enhanced microbial bile acid transformation. Administration of GR-7 effectively suppressed secondary bile acid accumulation in both early and late-stage interventions (Figs. 3c–d, all *P* < 0.05). Notably, low-dose GR-7 was sufficient to reduce total secondary bile acids, whereas high-dose treatment exerted a more pronounced effect on total secondary unconjugated bile acids (Fig. 3c), further supporting dose-dependent suppression of microbial bile acid deconjugation and downstream secondary bile acid formation. Collectively, these findings demonstrate that GR-7 robustly changes cecal bile acids through inhibition of bacterial BSH activity in vivo.

**Figure 3.**
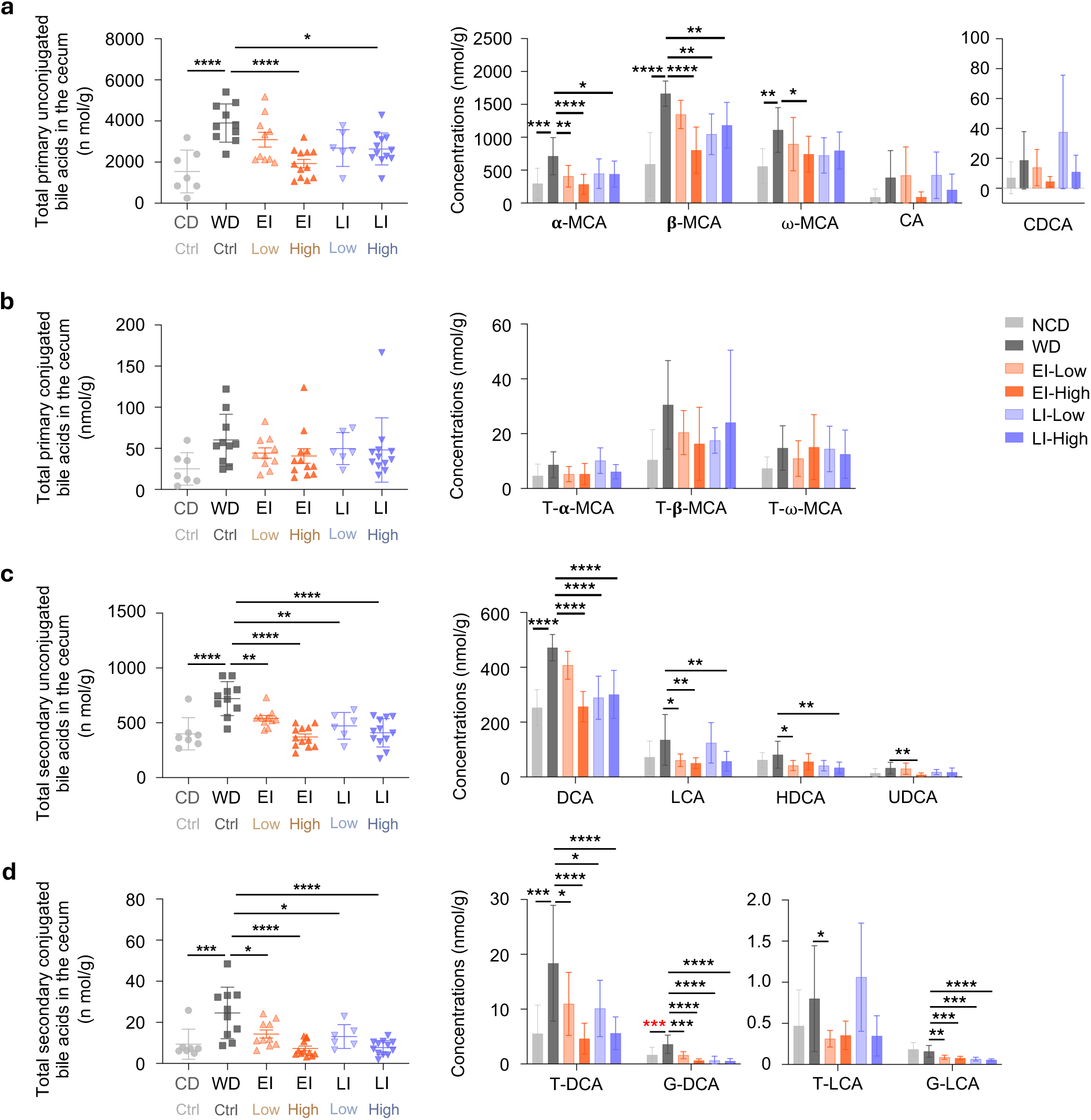
GR-7 changes cecal bile acid composition. **a**, Total primary unconjugated bile acids in cecal contents (left) and concentrations of representative individual primary unconjugated bile acid species, including α-muricholic acid (α-MCA), β-muricholic acid (β-MCA), ω-muricholic acid (ω-MCA), cholic acid (CA), and chenodeoxycholic acid (CDCA) (right). **b**, Total primary conjugated bile acids in cecal contents (left) and concentrations of representative individual taurine-conjugated primary bile acids, including tauro-α-muricholic acid (T-α-MCA), tauro-β-muricholic acid (T-β-MCA), and tauro-ω-muricholic acid (T-ω-MCA) (right). **c**, Total secondary unconjugated bile acids in cecal contents (left) and concentrations of representative individual secondary unconjugated bile acid species, including deoxycholic acid (DCA), lithocholic acid (LCA), hyodeoxycholic acid (HDCA), and ursodeoxycholic acid (UDCA) (right). **d**, Total secondary conjugated bile acids in cecal contents (left) and concentrations of representative individual taurine- or glycine-conjugated secondary bile acids, including tauro-deoxycholic acid (T-DCA), glyco-deoxycholic acid (G-DCA), tauro-lithocholic acid (T-LCA), and glyco-lithocholic acid (G-LCA) (right). Cecal bile acids were quantified by targeted bile acid profiling. Data are shown as mean ± SD. Each symbol represents one mouse. Statistical significance was determined by one-way ANOVA with Bonferroni’s multiple-comparisons test. * *P* < 0.05; ** *P* < 0.01; *** *P* < 0.001; **** *P* < 0.0001.

### BSH inhibition reshapes hepatic gene expression in MASH during late intervention

To further elucidate the molecular mechanisms underlying the therapeutic effects of BSH inhibition, RNA sequencing was performed on liver tissues. Because late intervention with high-dose GR-7 was most effective in suppressing MASH, subsequent transcriptomic analyses focused on comparisons between WD-fed mice and WD-fed mice receiving late intervention with high-dose GR-7. Substantial transcriptional reprogramming was observed in the liver following high-dose GR-7 treatment, with clear separation from WD-fed controls (Fig. 4a). KEGG pathway enrichment analysis revealed coordinated regulation of multiple metabolic and inflammatory pathways (Fig. 4b). Specifically, GR-7 administration significantly upregulated bile acid–related pathways, including steroid hormone biosynthesis, cholesterol metabolism, and bile secretion. In parallel, lipid metabolic pathways such as linoleic acid metabolism and arachidonic acid metabolism were also significantly modulated. In contrast, pathways associated with insulin resistance and inflammation—including cytokine–cytokine receptor interaction and TNF signaling—were markedly downregulated. These transcriptional changes were consistent with the observed phenotypic improvements in hepatic steatosis and inflammation following high-dose therapeutic BSH inhibition.

**Figure 4.**
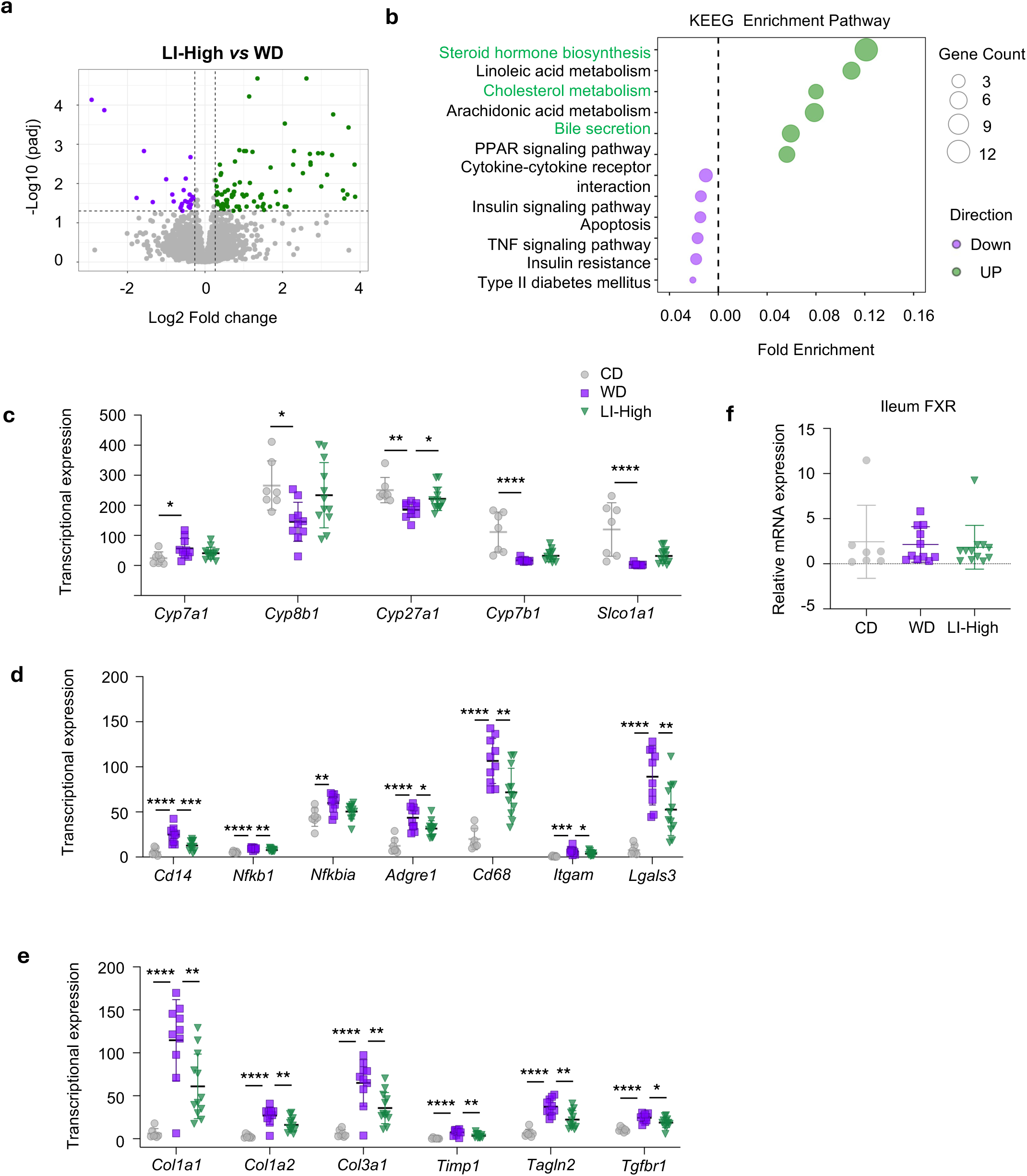
Late intervention with high-dose GR-7 reprograms hepatic transcriptome toward alternative bile acid synthesis and suppresses inflammatory and fibrotic signaling. **a**, Volcano plot showing differentially expressed hepatic genes in the LI-High group compared with WD controls. Dotted vertical lines indicate log₂ fold-change thresholds, and the horizontal dashed line indicates the adjusted *P*-value cutoff. Upregulated and downregulated genes are highlighted. **b,** KEGG pathway enrichment analysis of differentially expressed genes in livers from LI-High versus WD groups. Bubble size indicates gene count, and color denotes the direction of regulation (upregulated or downregulated). Differentially expressed genes were defined using an adjusted *P*-value cutoff of 0.1 and an absolute fold-change threshold of 1.2. **c**, Hepatic expression of bile acid synthesis and transport–related genes, including *Cyp7a1*, *Cyp8b1*, *Cyp27a1*, *Cyp7b1*, and *Slco1a1*. **d**, Hepatic expression of inflammation- and immune-related genes, including *Cd14*, *Nfkb1*, *Nfkbia*, *Adgre1*, *Cd68*, *Itgam*, and *Lgals3*. **e**, Hepatic expression of fibrosis-related genes, including *Col1a1*, *Col1a2*, *Col3a1*, *Timp1*, *Tagln2*, and *Tgfbr1*. **f**, Relative ileal *Fxr* mRNA expression determined by quantitative RT-PCR (qPCR). **c-e,** Gene expression levels were quantified by RNA sequencing and are shown as normalized transcript abundance. CD: chow diet; WD: Western diet; LI-High: Late intervention with high dose. Data are shown as mean ± SD. Each symbol represents one mouse. Statistical significance was assessed using the one-way ANOVA with Bonferroni’s multiple comparisons test. * *P* < 0.05; ** *P* < 0.01; *** *P* < 0.001; **** *P* < 0.0001.

To further dissect bile acid–specific regulatory mechanisms, we examined the expression of genes involved in hepatic bile acid synthesis (Fig. 4c). Compared with chow-fed controls, WD feeding significantly increased expression of *Cyp7a1*, the rate-limiting enzyme of the classical bile acid synthesis pathway; this induction was not significantly altered by BSH inhibitor treatment. In contrast, WD feeding suppressed *Cyp27a1*, a key enzyme in the alternative bile acid synthesis pathway, and this suppression was robustly reversed by GR-7, indicating a shift toward enhanced alternative bile acid synthesis. Similarly, downstream enzymes involved in bile acid synthesis, including *Cyp8b1* and *Cyp7b1*, were downregulated under WD conditions and showed a trend toward restoration following BSH inhibitor treatment, although these changes did not reach statistical significance. In addition to bile acid synthesis, hepatic bile acid transport was also affected. Among bile acid transporters, *Slco1a1* was significantly downregulated by WD feeding but exhibited a trend toward restoration in the BSH inhibitor–treated group, suggesting enhanced hepatocellular uptake of portal bile acids in response to intestinal bile acid remodeling.

Analysis of inflammatory gene expression further supported an anti-inflammatory effect of BSH inhibition. Multiple genes upregulated by WD feeding were significantly suppressed by GR-7 treatment, including *Cd14* and *Nfkb1*, while *Nfkbia* showed a trend toward reduction (Fig. 4d). These genes are associated with gut barrier dysfunction and inflammatory signaling. Moreover, macrophage- and Kupffer cell–associated markers such as *Adgre1*, *Cd68*, *Itgam*, and *Lgals3* were also markedly reduced, indicating attenuation of hepatic innate immune activation (Fig. 4d). Although late-stage intervention with high-dose GR-7 treatment did not result in statistically significant improvement in histological fibrosis, transcriptomic analysis revealed significant downregulation of several fibrosis-associated genes, including *Col1a1, Col1a2, Col1a3, Timp1, Tagln2, and Tgfbr1* (Fig. 4e). In addition, quantitative PCR analysis of ileal FXR expression showed no significant difference following high-dose GR-7 treatment (Fig. 4f). These changes indicate that BSH inhibition may exert early anti-fibrotic effects at the transcriptional level that precede detectable histological improvement.

Taken together, these transcriptomic data demonstrate that late therapeutic intervention with high-dose BSH inhibitor induces broad transcriptional reprogramming in the liver, characterized by enhanced bile acid metabolic remodeling, suppression of inflammatory signaling, and attenuation of fibrosis-related gene expression, thereby contributing to the overall improvement of MASH pathology.

### CyTOF profiling reveals immune remodeling induced by western diet and late intervention with BSH inhibitor

To comprehensively characterize hepatic immune alterations induced by WD feeding and late intervention with GR-7, we performed high-dimensional mass cytometry (CyTOF) analysis on liver immune cells (Fig. 5a). Unsupervised CyTOF analysis demonstrated that, compared with chow diet–fed mice, WD feeding profoundly remodeled the hepatic immune compartment. Specifically, WD-fed mice exhibited a significant reduction in T cells accompanied by an increase in B cells (Fig. 5b). Overall CD4 T cells were significantly decreased, whereas regulatory CD4 T cells (Tregs) were increased (Fig. 5c). At the functional level, CyTOF profiling further revealed significant suppression of IL-17A⁺ CD4 T cells and granzyme B⁺ CD8 T cells (Fig. 5d). Collectively, these data indicate that western diet feeding drives the liver into an immunosuppressed yet dysregulated state, characterized by impaired adaptive immune effector functions and compensatory expansion of regulatory immune populations, rather than overt inflammatory activation.

**Figure 5.**
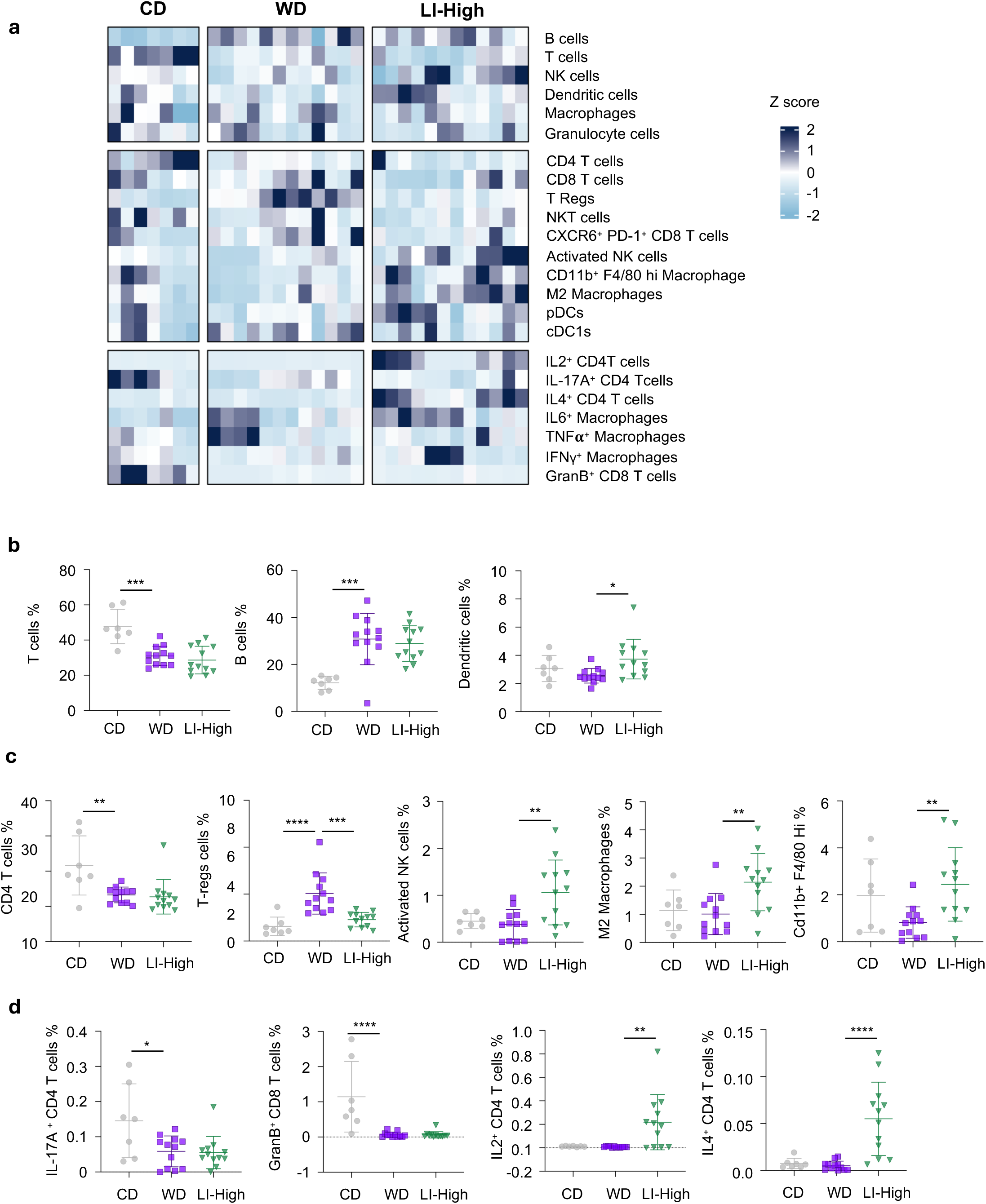
Late high-dose GR-7 remodels the hepatic immune landscape in western diet–fed mice. **a**, Heatmap summarizing CyTOF-based profiling of major hepatic immune cell populations, functional subpopulations, and cytokine-producing subsets in chow diet (CD), western diet (WD), and late-intervention high-dose GR-7 (LI-High) groups. Data are shown as Z scores for each immune population across individual mice. **b**, Percentages of total T cells, B cells, and dendritic cells among live hepatic immune cells. **c**, Percentages of immune cell subpopulations, including CD4 T cells, regulatory T cells (Tregs), activated NK cells, M2 macrophages, and CD11b⁺F4/80^hi^ macrophages. **d**, Percentages of cytokine-producing immune cell subsets, including IL-17A⁺ CD4 T cells, Granzyme B⁺ CD8 T cells, IL-2⁺ CD4 T cells, and IL-4⁺ CD4 T cells. CyTOF analysis was performed on liver immune cells isolated at the study endpoint. Data are shown as mean ± SD. Each symbol represents one mouse. Statistical significance was determined by one-way ANOVA with Bonferroni’s multiple comparisons test. * *P* < 0.05; ** *P* < 0.01; *** *P* < 0.001; **** *P* < 0.0001.

**Figure 6.**
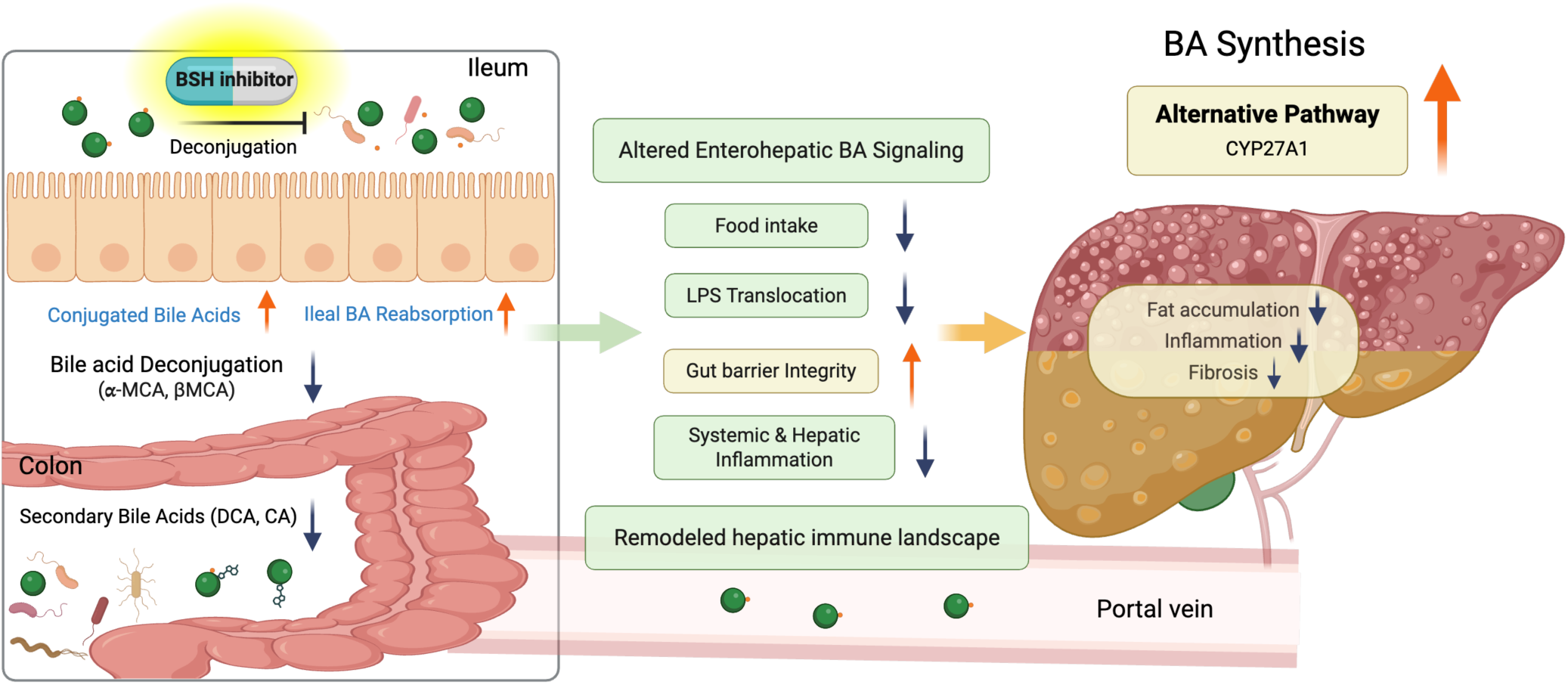
Mechanistic model illustrating how BSH inhibition remodels enterohepatic bile acid signaling and attenuates MASH progression. Pharmacological inhibition of microbial bile salt hydrolase (BSH) suppresses intestinal bile acid deconjugation, leading to increased levels of conjugated bile acids (e.g., TβMCA) and enhanced ileal bile acid reabsorption. Reduced deconjugation limits microbial conversion of primary bile acids into cytotoxic secondary bile acids, including deoxycholic acid (DCA) and lithocholic acid (LCA), thereby decreasing the intestinal and systemic secondary bile acid pool. These changes improve intestinal barrier integrity, reduce microbial lipopolysaccharide (LPS) translocation, and attenuate systemic and hepatic inflammation. In the liver, altered bile acid signaling promotes a shift from the classical to the alternative bile acid synthesis pathway (CYP27A1), resulting in reprogrammed hepatic bile acid metabolism with reduced secondary bile acid toxicity. Collectively, these coordinated effects contribute to decreased hepatic steatosis, inflammation, and fibrosis, thereby ameliorating MASH pathology.

CyTOF-based immune profiling further revealed that late intervention with high-dose BSH inhibitor did not globally normalize hepatic immune cell distributions to chow-fed levels; instead, several immune populations were selectively altered compared with WD-fed mice. Notably, dendritic cells were significantly increased following late intervention with high-dose treatment, indicating alterations in antigen-presenting cell compartments during late-stage intervention (Fig. 5b). Among immune subpopulations, late intervention with high-dose treatment resulted in a significant reduction in Tregs compared with WD-fed mice. In parallel, CyTOF analysis demonstrated a significant increase in activated NK cells, together with an increase in M2 macrophages and CD11b⁺ F4/80^hi^ macrophages, indicating altered innate immune activation and macrophage polarization during late-stage intervention (Fig. 5c). At the T cell cytokine level, late intervention with high-dose treatment significantly increased IL-2⁺ CD4 T cells and IL-4⁺ CD4 T cells, consistent with enhanced immunoregulatory and Th2-associated cytokine programs (Fig. 5d). Taken together, these results demonstrate that late intervention with high-dose BSH inhibition does not fully restore hepatic immune composition to a chow-fed state, but instead selectively reshapes the immune microenvironment by recalibrating regulatory, innate, and cytokine-associated immune programs during established disease.

## Discussion

Here, we demonstrate that pharmacological inhibition of microbial BSH using the inhibitor GR-7 exerts stage- and dose-dependent therapeutic effects in a mouse model of MASH. GR-7 reduced food intake and body weight gain. Early intervention with GR-7 robustly suppressed hepatic fibrosis even at a low dose. Although early and late intervention reduced hepatic steatosis and injury, only late-stage administration of high-dose GR-7 exerted pronounced effects on hepatic steatosis, inflammation, and injury. However, fibrosis was not histologically reversed at this stage and was instead suppressed primarily at the transcriptional level. Collectively, these findings indicate that microbial BSH inhibition holds substantial therapeutic potential for MASH, with efficacy strongly influenced by both treatment timing and dosage.

We further validated the in vivo efficacy of GR-7 as a BSH inhibitor by profiling bile acid composition in the cecum, a major site of microbial bile acid metabolism. High-dose GR- 7 significantly reduced total unconjugated primary bile acids in both early and late intervention settings, indicating that sufficient compound exposure is required to effectively engage the catalytic cysteine residue of BSH. Notably, the changes observed in the primary bile acid pool following GR-7 treatment were driven predominantly by MCA species, which constitute major primary bile acids in rodents. In contrast, CA, a predominant primary bile acid in humans but only a minor component of the rodent bile acid pool, also exhibited a reduction trend following GR-7 treatment, suggesting that the effects of GR-7 are not exclusively limited to rodent-specific bile acid species. This observation is consistent with previous reports demonstrating that BSH inhibitors exhibit broad-spectrum inhibitory activity across diverse bacterial strains and achieve effective target engagement in vivo.^15^ Remarkably, GR-7 also reduced secondary bile acids, including both conjugated and unconjugated species, consistent with suppressed microbial bile acid deconjugation and downstream secondary bile acid formation. Although it remains unclear whether this effect reflects direct inhibition of secondary bile acid–producing pathways or indirect changes in microbial composition, the pronounced suppression of the highly hydrophobic bile acids deoxycholic acid (DCA) and lithocholic acid (LCA) is particularly relevant. Elevated levels of DCA and LCA have been implicated in hepatocyte injury, intestinal barrier dysfunction, and inflammatory signaling, and metabolomic profiling linked increased DCA and LCA levels with advanced liver fibrosis in patients.^9,17^ These findings align with our observations, as GR-7 administration reduced cecal DCA and LCA levels and was associated with improved hepatic fibrosis. Mechanistically, these effects may be mediated, at least in part, through improved intestinal barrier integrity. Consistent with this hypothesis, prior studies have shown that BSH inhibition protects gut epithelial monolayers from damage induced by microbial-derived unconjugated bile acids via altered micelle formation in vitro.^15^ In the present study, we observed similar protective effects in vivo, as evidenced by significantly reduced plasma LPS levels in the late intervention group receiving high-dose GR-7, with a similar downward trend observed in the early intervention group. Collectively, these findings indicate that high-dose GR-7 effectively reduces cecal unconjugated primary and secondary bile acids and confers hepatoprotective effects by attenuating hepatic inflammation, at least in part via restoration of gut barrier integrity and reduced microbial translocation.

Given that GR-7 is not intestinally absorbed,^18^ our findings indicate that inhibition of microbial BSH can nonetheless reprogram hepatic bile acid metabolism through altered gut–liver signaling. Hepatic RNA-sequencing revealed upregulation of bile acid–related pathways, accompanied by increased expression of *Cyp27a1*, indicating activation of the alternative bile acid synthesis pathway. Consistent with our findings, recent studies have demonstrated that inhibition of microbial BSH activity can activate the alternative bile acid synthesis pathway and confer beneficial metabolic effects. For example, administration of theabrownin, a key bioactive component of Pu-erh tea, suppresses intestinal bacterial BSH activity, promotes alternative bile acid synthesis, and reduces cholesterol and lipid levels.^19^ In addition, other naturally occurring BSH inhibitors, including riboflavin and grape seed extracts, have also been reported to activate the alternative bile acid synthesis pathway, suggesting a conserved metabolic response to BSH inhibition.^20,21^ Mechanistically, suppression of microbial bile acid deconjugation is expected to increase the intestinal pool of conjugated bile acids, which are weaker FXR agonists, thereby attenuating ileal FXR signaling and relieving FXR–FGF15–mediated feedback inhibition of hepatic bile acid synthesis. This shift favors redirection from the classical CYP7A1-dependent pathway toward the alternative CYP27A1-mediated pathway, promoting cholesterol utilization and the production of more hydrophilic bile acids.^22^ Although we did not observe changes in ileal FXR expression, the dynamic nature of FXR signaling and rapid enterohepatic bile acid cycling may limit detection of robust transcriptional differences in vivo. Notably, activation of the alternative bile acid synthesis pathway has been proposed as a compensatory, protective response in advanced MASLD and fibrosis,^23^ which may explain the greater therapeutic efficacy of GR-7 during late intervention compared with early intervention, highlighting disease stage–dependent metabolic plasticity as a key determinant of responsiveness to microbial BSH inhibition.

Furthermore, high-dose early intervention was associated with a significant reduction in food intake, which was also validated in wild-type mice. This effect may have partially contributed to the attenuation of MASLD progression, particularly with respect to hepatic lipid accumulation and body weight gain. Consistent with previous reports, mono-colonization with bile salt hydrolase–deficient *Bacteroides thetaiotaomicron* reduced body weight and food consumption through altered bile acid signaling rather than impaired nutrient absorption.^11^ This metabolic reprogramming was accompanied by enhanced glycolysis and increased lipid uptake in ileal epithelial cells, supporting coordinated lipid utilization and energy homeostasis.^11^

Lastly, we investigated whether inhibition of microbial bile salt hydrolase (BSH) alters the hepatic immune landscape in MASH. Unbiased high-dimensional immune profiling by CyTOF revealed that western diet feeding induced profound remodeling of the hepatic immune compartment. However, GR-7 treatment did not globally restore these diet-induced immune alterations. Instead, it selectively reshaped the hepatic immune microenvironment by recalibrating regulatory, innate, and cytokine-associated immune programs during established disease. Notably, late intervention with high-dose GR-7 was associated with a shift in hepatic macrophage polarization toward an M2-like, immunoregulatory phenotype. This macrophage remodeling was accompanied by coordinated downregulation of inflammatory and macrophage-associated genes, including *Adgre1*, *Cd68*, *Itgam*, and *Lgals3*, which are closely linked to Kupffer cell activation. Together, these findings suggest that GR-7 promotes a more immunoregulatory hepatic environment rather than broadly suppressing immune responses, consistent with an improved inflammatory state during late-stage MASH. These immune changes are mechanistically consistent with GR-7–mediated reshaping of intestinal bile acid composition, particularly through reduction of hydrophobic and cytotoxic secondary bile acids such as DCA and LCA. Reduced portal endotoxemia would be expected to attenuate Toll-like receptor–mediated activation of hepatic macrophages, thereby limiting inflammatory Kupffer cell signaling and favoring a macrophage-dominant immunoregulatory state in established MASH.

In summary, our study demonstrates that pharmacological inhibition of microbial BSH by the non-absorbable compound GR-7 exerts stage- and dose-dependent beneficial therapeutic effects in a mouse model of MASH. By suppressing microbial bile acid deconjugation, GR-7 reshapes the intestinal bile acid pool, reduces hydrophobic and cytotoxic secondary bile acids such as DCA and LCA, and improves intestinal barrier function, thereby limiting portal endotoxemia. These changes reprogram hepatic bile acid metabolism toward activation of the alternative CYP27A1-mediated synthesis pathway, promote cholesterol utilization, and alleviate metabolic and inflammatory stress, particularly during late-stage disease. In addition, reduced food intake might have partially contributed to the metabolic benefits of GR-7. Together, these findings establish microbial BSH inhibition as a promising microbiome-targeted therapeutic strategy for MASH.

## Acknowledgments

This study was supported by a lab service agreement with Vertero Therapeutics (formerly Axial Therapeutics) and services provided by NIH center P30 DK120515. We thank Dr. Sloan Devlin for helpful discussions related to this work.

## Author contributions

W.W. performed animal experiments and biochemical analyses and drafted the manuscript. R.G. designed the study, oversaw study execution, including Cytof, RNAseq, and bile acid analyses, and analyzed data. Y.W. performed animal experiments. C.J.O. and J.T.L. oversaw study design and execution, C.J.O. oversaw the production of GR-7. X.W. assisted with RNA-seq analysis. M.C. and J.S. designed and oversaw Cytof analyses. M.C.C. contributed to study design. S.G. was responsible for breeding and genotyping of mice. T. Y. assisted with animal experiments. A.H., S.C. and M.F. assisted with mouse harvesting and sample collection. D.D. assisted with mouse breeding and genotyping. A.S.C, R.K.S, and B.S. were responsible for study concept and design, interpretation of data, editing the manuscript, and study supervision. All authors read and edited the manuscript.

